# Conformational clamping by a membrane ligand activates the EphA2 receptor

**DOI:** 10.1101/2021.04.08.439029

**Authors:** Justin M. Westerfield, Amita R. Sahoo, Daiane S. Alves, Brayan Grau, Alayna Cameron, Mikayla Maxwell, Jennifer A. Schuster, Paulo C. T. Souza, Ismael Mingarro, Matthias Buck, Francisco N. Barrera

## Abstract

The EphA2 receptor is a promising drug target for cancer treatment, since EphA2 activation can inhibit metastasis and tumor progression. It has been recently described that the TYPE7 peptide activates EphA2 using a novel mechanism that involves binding to the single transmembrane domain of the receptor. TYPE7 is a conditional transmembrane (TM) ligand, which only inserts into membranes at neutral pH in the presence of the TM region of EphA2. However, how membrane interactions can activate EphA2 is not known. We systematically altered the sequence of TYPE7 to identify the binding motif used to activate EphA2. With the resulting six peptides, we performed biophysical and cell migration assays that identified a new potent peptide variant. We also performed a mutational screen that determined the helical interface that mediates dimerization of the TM domain of EphA2 in cells. These results, together with molecular dynamic simulations, allowed to elucidate the molecular mechanism that TYPE7 uses to activate EphA2, where the membrane peptide acts as a molecular clamp that wraps around the TM dimer of the receptor. We propose that this binding mode stabilizes the active conformation of EphA2. Our data, additionally, provide clues into the properties that TM ligands need to have in order to achieve activation of membrane receptors.

## Introduction

EphA2 is a receptor tyrosine kinase (RTK) that directs blood vessel remodeling and nervous system formation during embryogenesis [1]. However, EphA2 can also drive oncogenesis and malignancy in several cancer types, and its expression is correlated with poor prognosis and metastasis [2]. The reason EphA2 plays such roles is that its activation causes remodeling of the cytoskeleton and affects cell adhesion, proliferation and differentiation [3].

EphA2 is activated by the ephrin ligands, which are membrane-anchored proteins found at the membrane of nearby cells. Specifically, ephrinA1 and ephrinA3 [4, 5], are anchored to the cell surface by a glycosylphosphatidylinositol (GPI) group. The interaction between EphA2 and ephrins ensures that initial contacts are established between the cells that bear each protein. When this contact is lost, for instance by down-regulation of ephrinA1, the signaling mechanism changes to one promoting cell migration and detachment, inducing cancer metastasis [1].

EphrinA1 binds to EphA2 at the ligand-binding domain (LBD) present at the extra-cellular region (ECR). This binding event causes a rearrangement of EphA2 that involves dimerization and other conformational changes. Multiple segments of the EphA2 receptor participate in receptor self-assembly: the LBD and the Cys-rich domain at the ECR [6, 7], the transmembrane (TM) domain, and the intra-cellular region (ICR). EphA2 dimerization leads to trans-activation of the kinase domain located at the ICR, which phosphorylates cytoplasmic tyrosine residues, triggering a signaling cascade. This ligand-dependent conformational rearrangement “ flows” from the LBD and the rest of the ECR into the plasma membrane through the single TM domain of EphA2 and the nearby juxtamembrane (JM) segment, and finally into the ICR kinase domain.

We recently reported that EphA2 can be activated by an alternative mechanism. Specifically, we designed TYPE7, a synthetic ligand that binds to the membrane region of EphA2 [8]. TYPE7 is a peptide that comprises the TM region of EphA2 modified by the introduction of glutamic acid (E) residues at strategic positions (**Fig. 1**). These E residues alter the overall hydrophobic nature of the EphA2 TM stretch and endow this sequence with behavioral flexibility. Indeed, TYPE7 is characterized by high solubility in physiological buffers, where it loses the characteristic α-helical nature of the TM to become unstructured. However, in these conditions TYPE7 is still able to bind to lipid membranes, albeit weakly and still in an unstructured conformation [8]. This process can be reversed by a pH drop, which protonates the E residues. The resulting loss of negative charge causes TYPE7 to transition from the membrane surface to its interior into a helical TM conformation. TYPE7 is therefore a conditional TM domain that is controlled by the environmental pH (8).

**Figure 1.**
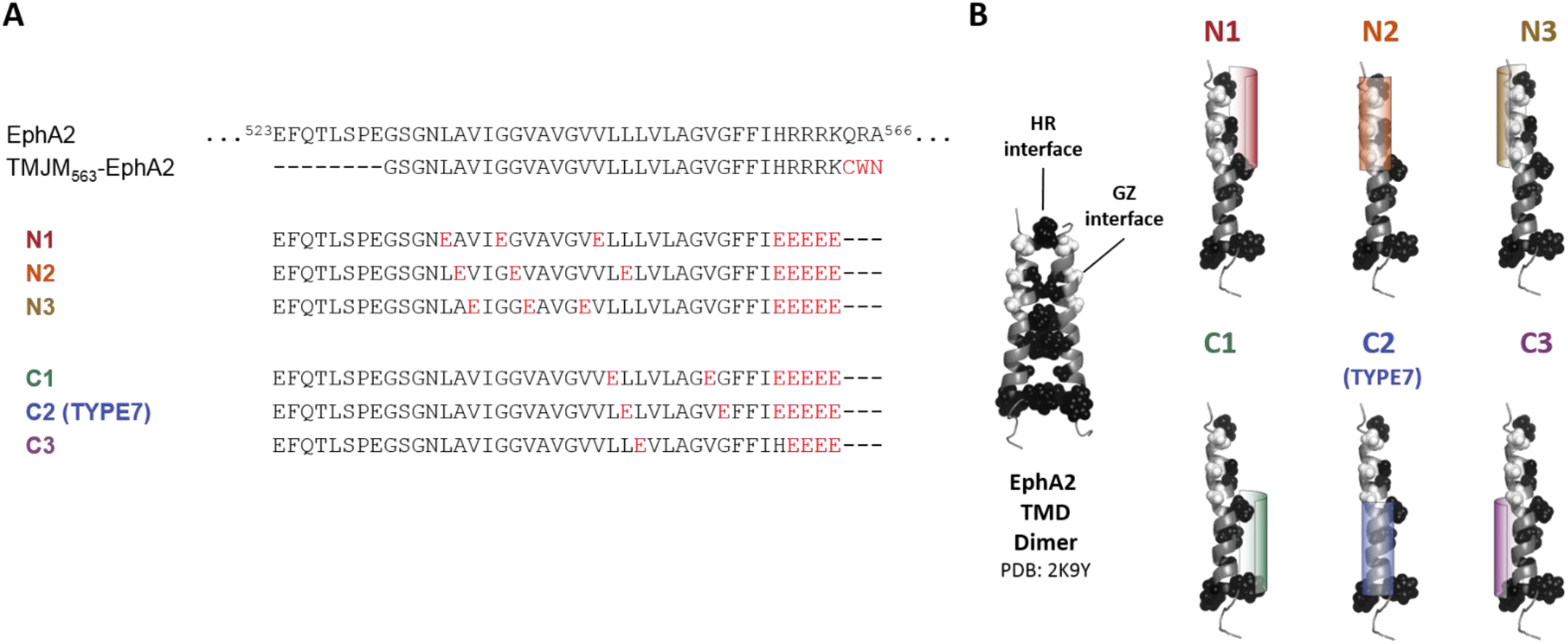
TYPE peptide sequences and design principle. **(A)** Sequences employed, where non-native residues are shown red. A CWN tag was added to the end of TMJM-EphA2_563_ for quantification purposes. **(B)** The NMR structure of the TM dimer of EphA2 is shown with HR residues shown in black, and GZ interfacial residues in white. Color shades mark the relative positions of the E “ shield” in the different α-helical TYPE peptides. Peptides were named for the interhelical interface that the E residues were added to: “ N” and “ C” denote vertical positions of E residues. Numbers (1, 2 & 3) represent rotational positions. EphA2 sequence information was taken from UniProt entry P29317-1.

TYPE7 is an efficacious ligand of EphA2, as it activates its kinase activity and causes trans-phosphorylation of Tyr 772. These events lead to inhibition of cell migration by de-phosphorylation of Akt and the resulting signaling cascade [8]. Interestingly, these effects are similar to those caused by ephrinA1 binding. However, it is not understood how TYPE7 activates EphA2. We previously showed that TYPE7 can interact with the TM and JM regions of EphA2 [8]. Developing a mechanistic understanding of how a membrane ligand activates EphA2 first requires revealing the molecular interface that EphA2 and TYPE7 use to interact in the membrane, and identifying the residues that participate in the formation of the complex. This information is required to rationalize how an interaction at the TM domain of EphA2 can largely recapitulate the receptor conformational changes caused by ephrinA1 docking at the LBD.

To understand how TYPE7 binds to the membrane region of EphA2, we systematically rotated the E residues characteristic of TYPE7 around the helical axis (**Fig. 1B**). We performed biophysical and cell biology experiments with the resulting six peptides, which provided insights into the binding motif that TYPE7 uses to bind to and activate the EphA2 receptor. Molecular dynamics simulations of the complex formed between the membrane region of EphA2 and an activating peptide reveal the molecular interactions that help to stabilize the EphA2 receptor in an active conformation.

## Results

We designed five variants of the TYPE7 peptide, where the key glutamic acid (E) residues were repositioned along the α-helix. These and the wild type peptide are expected to position each the glutamic acids at a unique TM helical face (**Fig. 1**). The rationale for adding E residues is that their side chains are bulky and are expected to create steric clashes that prevent TM-TM interactions exclusively at that interface. Additionally, E residues grant the peptides solubility in buffer, and their deprotonation at acidic pH triggers membrane insertion of TYPE7 [8]. The E residues were placed at three major helical interfaces. Interface 1 corresponds to the helix-helix contacts observed in the NMR structure of the TM EphA2 dimer [9]. This interface is mediated by a heptad repeat (HR) motif [4] and is proposed to be populated when EphA2 undergoes ligand-independent (non-canonical) activation [10]. Interface 2 is separated by a ∼140 degree rotation around the helical axis from interface 1. Interface 2 contains a Glycine Zipper (GZ) (**Fig. 1B**), a common motif in transmembrane dimer formation that contains a GxxxG motif, also known as GAS-right [11-14]. The GZ interface is not observed in the NMR structure, but it has been computationally predicted to mediate dimerization in the ligand-activated conformation of EphA2 [4]. Finally, the third interface was selected as a control, as no dimerization interactions have been observed. The six TYPE peptides we describe in this work are classified as N or C, as E residues were placed either in the amino-terminal (N_t_) or the carboxy-terminal (C_t_) halves of the TM helix, respectively. Each peptide is systematically dubbed for the region of the helix (N_t_ or C_t_) and the interface (1, 2, or 3) where the acidic residues are placed. TYPE7 will be also referred herein as TYPE-C2, as E residues are at the C_t_ of interface 2 (GZ).

### Acidity triggers the membrane helical formation in all TYPE peptides

In order for the TYPE peptides to interact with TM domain of EphA2, as intended, first they must adopt a helical conformation in the membrane. We determined the secondary structure of the peptides using circular dichroism (CD) spectroscopy (**Fig. S1**). We observed that all the peptides behaved similarly to TYPE-C2, which exhibits pH-triggered membrane insertion. TYPE-C2 is highly soluble in aqueous solution, where it is unstructured (state I) [8]. The peptide binds to POPC liposomes at neutral pH (state II) in a largely random-coil conformation. Finally, TYPE-C2 forms a transmembrane helix when the solution is acidic (state III) [8]. The results show that the changes in E position along the helix do not prevent any of the new peptides to populate the three aforementioned states. To study the pH-dependency of helix formation we performed pH titrations of the six peptides using CD. **Figs. S2-3** show that all peptides gain helical structure in lipid membranes at acidic pH, and that the value of the pH where 50% of the helix is formed (pH_50_) is 5.4-6.0. These results indicate that the initial design principle was successful, and all peptides maintain pH-triggered membrane insertion.

### TYPE peptides interact with the membrane region of EphA2

Once we determined that the new peptides behaved similarly to TYPE-C2, which forms a TM helix at acidic pH, we studied if they interacted with the TM region of EphA2. To this end we used a liposome reconstitution system we developed previously for TYPE7 [8]. In this system, we prepared proteo-liposomes that contain a peptide corresponding to the membrane region of EphA2. Specifically, we used residues 531 to 563 of EphA2, consisting of the TMD region plus five juxtamembrane (JM) residues, and a C_t_ CWN tag added for quantification purposes (**Fig. 1A**). The resulting sequence is referred to as TMJM_563_-EphA2. In the liposome assay we study if the presence of TMJM_563_-EphA2 alters the pH-dependency of the membrane insertion of the TYPE peptides. Differences between the pH midpoints (pH_50_) obtained in liposomes and proteo-liposomes would reveal an interaction between the TYPE peptide and the membrane region of EphA2. For example, we observed that the presence of TMJM_563_-EphA2 promoted the membrane insertion of TYPE-C2, as less acidity is needed for helix formation (**Fig. 2**). This result is likely explained because the TM state of TYPE-C2 has available a new partner (TMJM_563_-EphA2) within the membrane. This causes a larger stabilization of the TM conformation than in plain bilayers. To perform this assay, the TYPE peptides were previously labeled at the N_t_ with the solvatometric probe 7-nitrobenzoxydiazole (NBD) [15]. NBD dehydration resulting from membrane insertion causes an increase in fluorescence intensity, and a blue-shift in the spectral center of mass, a parameter similar to the spectral maximum that can be obtained with higher accuracy using Eq. 1 [15]. As a control, we also measured the NBD fluorescence changes at increasing POPC concentrations, which demonstrated that the six peptides efficiently bind to liposomes (**Fig. S4**). These results confirm that all peptides populate state II, and are thus competent to bind lipid vesicles.

**Figure 2.**
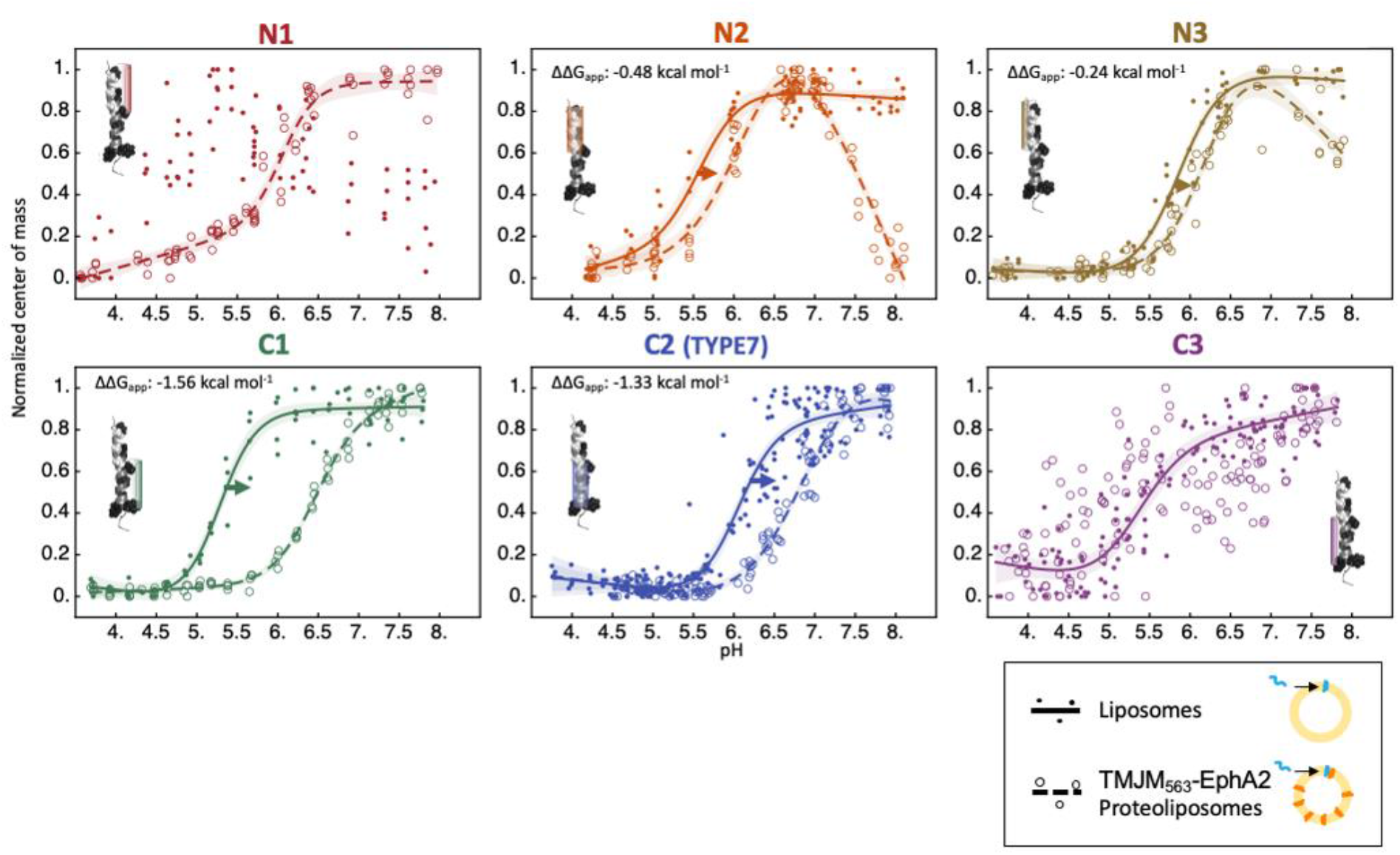
TYPE peptides interact with the TMD of EphA2. NBD-labeled TYPE peptides were incubated with POPC liposomes lacking (closed circles, solid lines) or containing (open circles, dashed lines) TMJM_563_EphA2. We measured the fluorescence center of mass as a function of pH. A blue-shifted (lower value) center of mass (Eq. 1) indicates burial of NBD in the bilayer. Fits were not performed if they did not show a clear sigmoidal shape (see panels N1 liposomes and C3 proteo-liposomes). Global fits are shown, and ± 95% confidence intervals are shaded. Means and fitting errors of the pH_50_ values derived from these fits are displayed in Table S1. The corresponding on-normalized can be found in **Fig. S5**.

**Fig. 2** shows that measuring NBD fluorescence changes as a function of pH resulted in sigmoidal membrane insertion curves in empty liposomes in all cases, except for TYPE-N1. Fitting the sigmoidal curves to eq. 2 provided the value of the pH_50_ for insertion (**Table S1)**. Interestingly, when experiments were repeated with proteo-liposomes, we observed that all TYPE peptides were affected by the presence of TMJM_563_-EphA2 in the vesicles (**Fig. 2, Table S1**), as the two curves did not overlay for any peptide. A shift to higher pH of the membrane insertion curve of TYPE-C2 was observed, as previously described [8]. For TYPE-C1, the changes were even larger, and the pH_50_ value increased by 1.2 pH units. More moderate increases occurred for the N2 and N3 peptides, which were also accompanied by the appearance of slopes at the baseline of the curve at neutral pH. The pH_50_ increase observed in these four peptides suggest that the presence of TMJM_563_-EphA2 promotes the formation of a TM helix, likely by establishing additional hydrophobic interactions that stabilize the TM conformation of the TYPE peptides. Finally, for the N1 and C3 peptides results were not conclusive, as a sigmoidal curve was only obtained either with proteoliposomes (N1) or “ non-doped” liposomes (C3). Similar conclusions were reached when changes in fluorescence intensity were measured instead to the spectral shift, although the data were noisier (**Fig. S6**). Overall, our results suggest that the C1 and C2 TYPE peptides interact with TMJM_563_-EphA2 more strongly than the other four TYPE peptides.

Taken together, our fluorescence results suggest that TMJM_563_-EphA2 interacts with the six TYPE peptides to different extents. The different “ morphologies” of the observed changes suggest that blocking TM interfaces with E residues can differently bias the conformation of the macromolecular complex formed in the membranes. Additionally, the observation that blocking of any single interface does not preclude interaction with TMJM_563_-EphA2, suggests that TYPE peptides use several interfaces to bind to the EphA2 receptor.

### Only two TYPE peptides (C1 and C2) inhibit cell migration

EphA2 activation by ephrinA1 results in a reduction of cell migration [16]. We previously demonstrated that the TYPE-C2 peptide specifically binds to EphA2 in the human malignant melanoma A375 cells [8]. This interaction activates the receptor to cause inhibition of EphA2-mediated cell migration. Similar results are obtained when cells are treated with a cross-linked version of the ephrinA1 ligand (Ea1), which recapitulates the membrane-clustered state of active ephrinA1. We performed a Boyden chamber assay to determine the effect of all TYPE variants on the migration of A375 cells in culture. **Fig. 3** shows that the C1 peptide inhibits cell migration similarly to C2 (TYPE7). These two peptides inhibit cell migration at least as efficiently as Ea1, suggesting that they achieve full inhibition of EphA2-driven cell migration. TYPE-C3 did not inhibit cell migration, while the smaller reductions observed for N1-3 were not statistically significant. The agreement we observed in the C1-2 peptides being the more efficient binders to EphA2 in the experiments shown in Figs. 2 and 3 indicate that these peptides are the best EphA2 ligands. These results also support the notion that that the proteo-liposome assay (**Fig. 2**) can provide physiologically relevant information.

**Figure 3.**
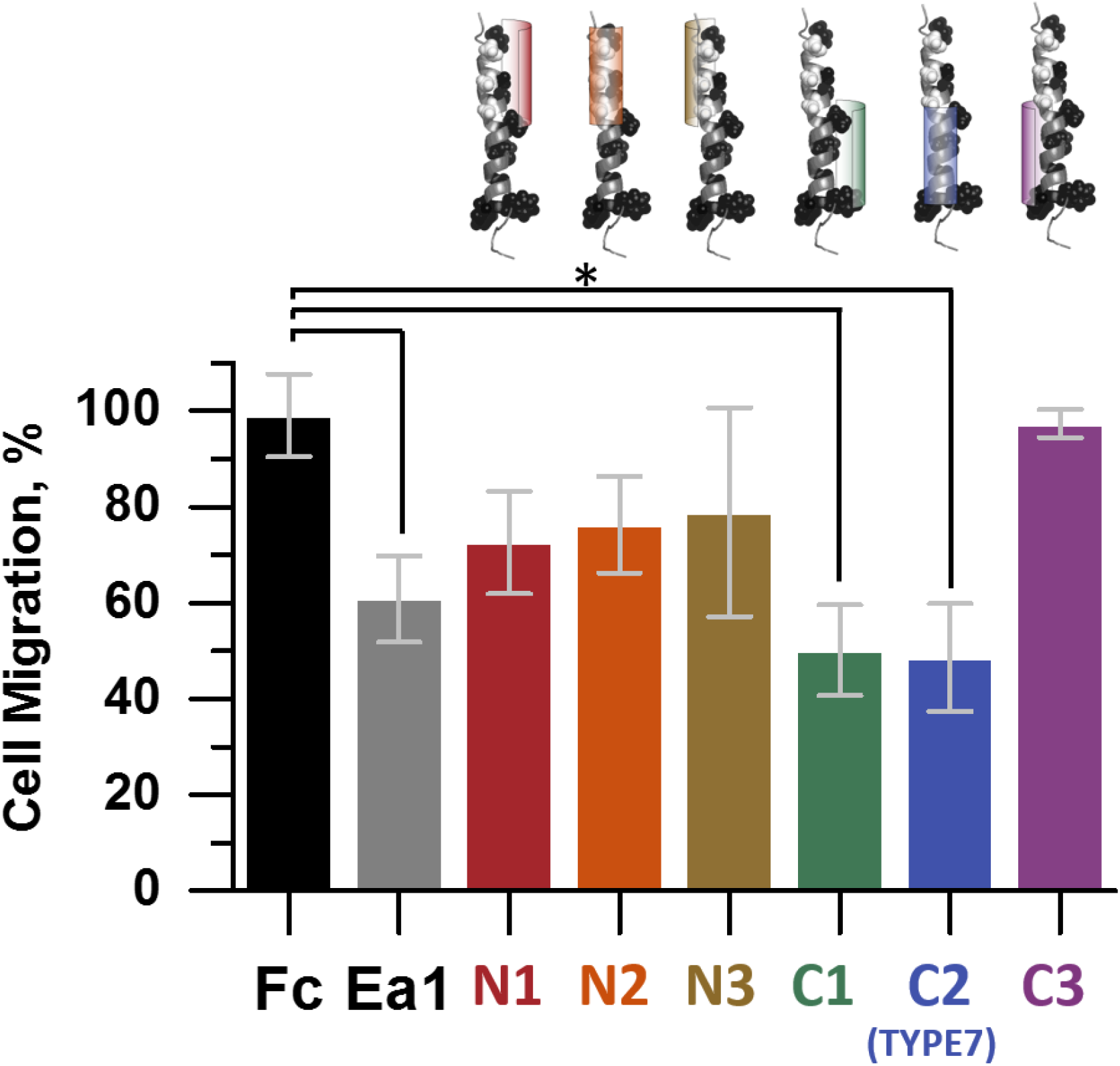
TYPE-C1 and TYPE-C2 efficiently reduce migration of A375 cells. Boyden chamber assay results measure cell migration. Data are normalized to control conditions. Inhibition of cell migration is achieved by incubation with saturating conditions of EphrinA1-Fc (Ea1). Fc is a negative control corresponding to the moiety present in Ea1. Averages ± standard deviation are shown. N ≥ 3. *, *p* < 0.05 when compared to Fc in a Dunnett’s T3 multiple means comparison test.

### The TMD of EphA2 dimerizes in cells using the GZ interface

Development of a model of the complex that EphA2 and TYPE-C2 form in the membrane first requires us to identify the helical interface/s of the EphA2 TMD dimer in the cell. However, the amino acids that participate in the interactions that allow the EphA2 TMD to dimerize have never been directly measured in cellular membranes. We performed a bimolecular fluorescence complementation (BiFC) assay for the TM region of EphA2, to directly identify the residues that mediate EphA2 dimerization [17]. We carried out a mutational analysis aimed at blocking TMD-TMD interactions at the three helical surfaces. To this end we replaced side chains at the different helical interfaces by the bulky side chain of Ile, to again create steric clashes that prevent dimerization [18]. Ile residues were used instead of Glu to ensure stable membrane insertion. Several mutations were performed at once, since single mutations in TM domains often have a small effect [19]. Specifically, 2 or 3 Ile residues were introduced at the GZ interface. We also replaced for Ile the HR positions mutated in the TYPE-N1 peptide. Finally, we mutated the residues at the third helical surface (same as in TYPE-N3), to cover the full helical rotation (**Fig. 4A**).

**Figure 4.**
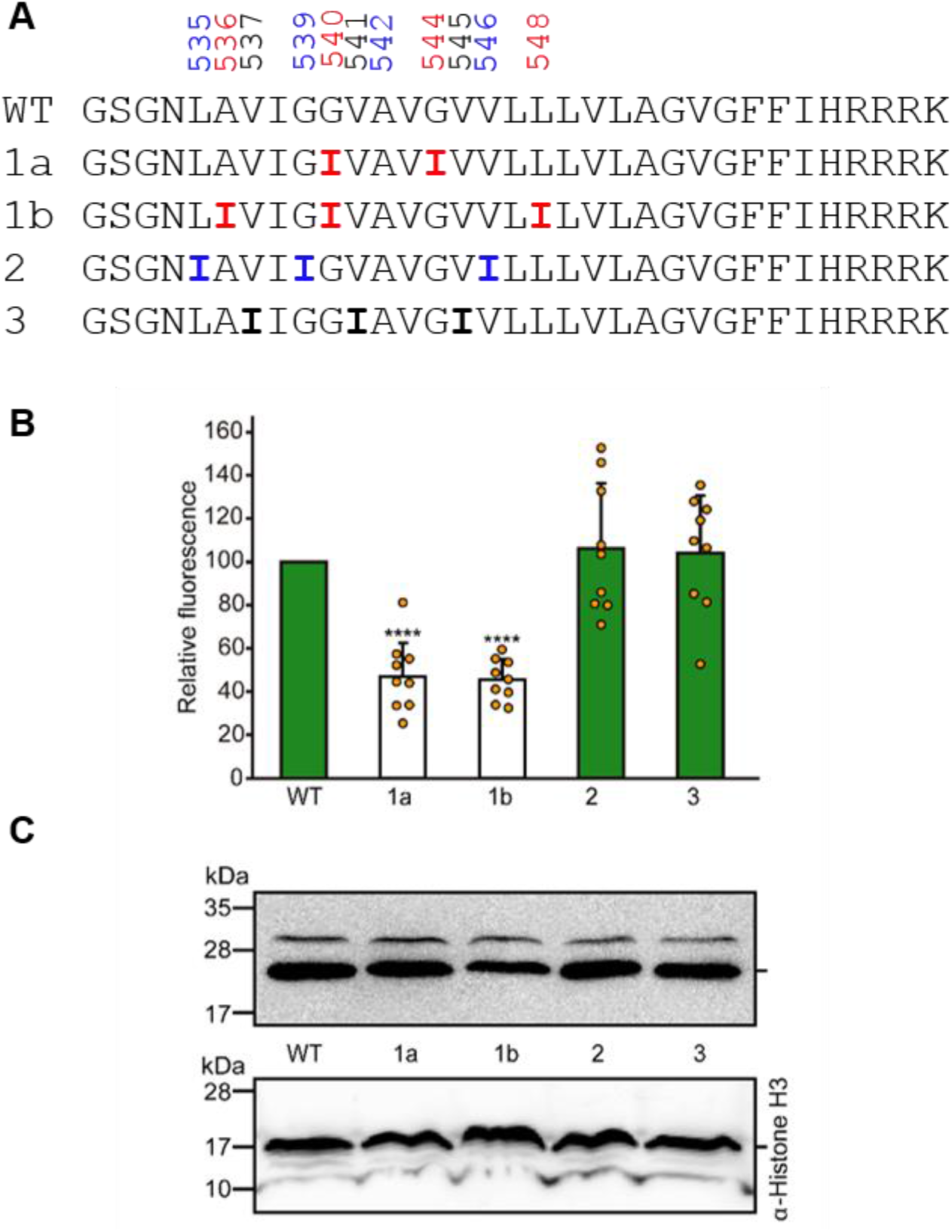
Measurement of self-assembly of EphA2 TMD in HEK293T cells. **(A)** Amino acid sequences fused to the VFP halves in the transfection plasmids are compared to the EphA2 WT sequence. Ile mutations were performed at interface 1, containing the GZ (red). GZ mutations are marked 1a and 1b. The other mutants were focused on interface 2 (HR, blue), and 3 (black). **(B)** VFP fluorescence is a reporter of self-assembly of the TM domain of EphA2. Error bars indicate standard deviation obtained from at least 9 independent experiments. EphA2 was set as reference value (*p* values of Student’s test: ****, < 0.0001). Green and white bars highlight the oligomerizing and non-oligomerizing TM segments, respectively. **(C)** Western blot showing the construct expression levels detected by α-c-Myc antibody. An α-Histone H3 antibody was used as a loading control.

To perform the BiFC experiment, we transfected HEK293T cells with two plasmids bearing the two halves of a split venus fluorescent protein (VFP), linked to the WT TMJM_563_-EphA2 sequence. Membrane dimerization of the TMD of EphA2 will bring together the two protein fragments, leading to VFP assembly and the resulting fluorescence emission [20]. Data in **Fig. 4B** show that the WT sequence efficiently brings together the two VFP halves, as expected in case of membrane dimerization. Mutation of the glycine residues in the helical interface 1, mediated by the GZ (G^540^I and G^544^I) led to a severe reduction in fluorescence. When we mutated other residues participating in the GZ interface (A^536^I, G^540^I and L^548^I), we obtained similar results. These data indicate that the GZ interface mediates dimerization of the isolated membrane region of EphA2 in the cells. Replacement of three key residues in the HR interface (L^535^I, G^539^I and V^546^I), on the other hand, did not alter VFP fluorescence. Similarly, no changes were observed for residues on the Val-rich interface 3 (V^537^I, V^541^I and V^545^I). These results suggest that a single helical face of EphA2 TMD, containing the GZ residues, mediates dimerization while the other two do not participate in the dimer formation in the membrane of human cells.

### Molecular dynamics (MD) simulations provide a mechanistic model for how the TYPE-C2 peptide activates EphA2

We want to understand how the TYPE-C2 peptide (TYPE7) and C1 can activate EphA2 and inhibit the metastatic phenotype characterized by cellular migration. This knowledge requires understanding the molecular interactions occurring between the two polypeptides when they form a hetero-complex in the membrane. Since obtaining high-resolution structures of membrane complexes in a lipid environment is currently challenging, we performed instead molecular dynamics simulations. These have the additional advantage that they can be carried out with TYPE peptides as monomers or as dimers, as the stoichiometry of the membrane complexes formed are not known.

We performed Coarse Grained (CG) MD simulations to study binding to TMJM_563_-EphA2 of the C2 peptide. We also used TYPE-N3 as a negative control. As starting points, the structure of a monomer of the C2 and N3 helices were modeled based on the NMR structure of the TM domain of EphA2, as they share most of the sequence [8]. The predicted monomer structures of the C2 and N3 peptides were used for subsequent CG simulation studies. First, we modeled the TMJM_563_-EphA2:C2 complex with a 2:1 stoichiometry, in such a way that both the TM monomers of EphA2 and the peptide were initially placed 50 Å apart from each other. The system was then subjected to a 4 μs CG simulation in a POPC bilayer. Each simulation was run in quadruplicate for both TMJM_563_-EphA2:C2 (2:1) and TMJM_563_-EphA2:N3 (2:1). We observed that the TMs of the EphA2 and the peptides efficiently interacted with each other (**Fig. S7**) forming a hetero-trimer. It should be noted that in all-atom (AT) MD simulations the configurational transitions in the lipid bilayer are typically slow, and we may not observe the peptides coming together in a native-like manner within several μs [21]. However, the energy landscape in the CG simulations is much smoother and diffusion is faster, allowing for an acceleration of the configurational and diffusional dynamics by a factor of up to 100-fold.

In order to study the association of the peptides, we monitored the helical distance between the two EphA2 TM monomers (named Chain A and Chain B henceforth) and the TYPE peptide (Chain C). We observed that the TYPE-C2/N3 peptides associated quickly in most of the cases with EphA2 TMs (Chains A and B) in the CG simulations (**Fig. S7**). **Fig. 5A** shows a superimposition of the central conformer of the major state of all 4 simulations of TMJM_563_- EphA2:C2. Interestingly, the association of the TYPE-C2 (chain C) allows the formation of the EphA2 TM dimer (chains A & B). In contrast, in the case of the TMJM_563_-EphA2:N3 complex (**Fig. 5B**), N3 breaks the association of EphA2. Control simulations of the isolated EphA2 TM showed that the dimers are formed early, at around 100 to 400 ns into the simulations, and were stable over 4 μs (**Fig. S8**). The results show that EphA2 forms a dimer with a different conformation than the NMR structure, in agreement with previous modeling [9].

**Figure 5.**
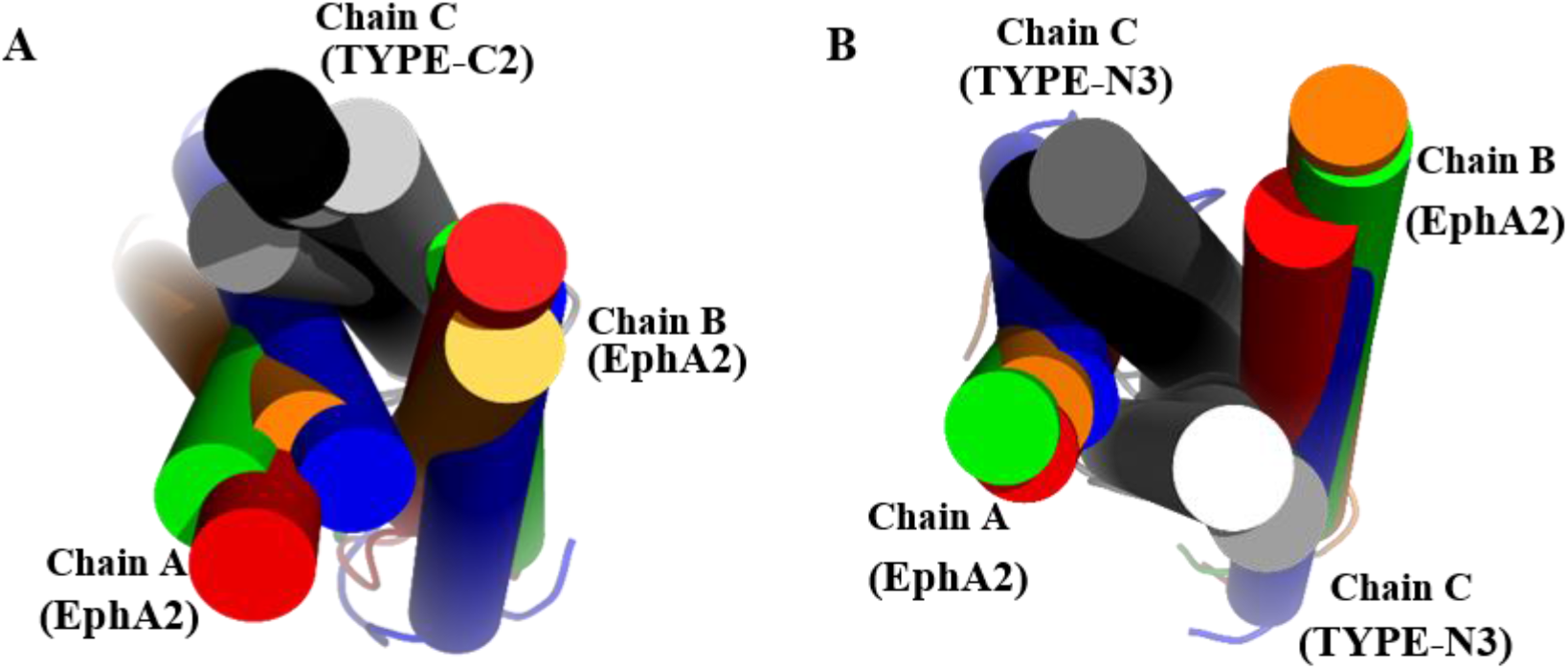
MD simulations show that TYPE-C2 and TYPE-N3 establish different interactions with EphA2. Superposition of the most highly populated conformations of the TM trimers (2:1 stoichiometry) corresponding to 4 independent simulations. Colored cylinders mark the different EphA2 helices (chain A & B) in different simulations, while chains C (TYPE peptides) are shown in white, grey, dark grey and black. **(A)** EphA2:C2 (2:1) system, in which the chain C (TYPE-C2) interacts with both chains A & B (TMs of EphA2), thereby facilitating their association towards active dimer conformation. **(B)** EphA2:N3 (2:1), where the chain C (N3) interacts differently with chains A and B forcing them to move outward, breaking their association. Similar results are obtained when the single C chain is “ above” or “ below” EphA2 (i.e., interposed from either side of the dimer).

**Table S2** shows the result of the PREDDIMER analysis of the EphA2 TM dimers (chain A & B) obtained for both the TYPE-C2/N3 systems, compared with the NMR structure of EphA2 TMDs and also with the control EphA2 simulation (**Fig. S8**). The dimers are characterized by the Fscor value -using the webserver PREDDIMER [22] in the analysis mode-as an indicator of packing tightness, and the helical crossing angle. For the NMR structure of the isolated EphA2 TM homodimer, the Fscor was 1.39, indicating a weak dimer (an Fscor of > 2.5 is regarded as a cut-off for predicting stable helix-helix interactions). The crossing angle is +23 degrees, corresponding to the left-handed dimer using the HR interface. For the control EphA2 system, the Fscor ranges from 2.1 to 3.6, suggesting a strong association between the TM monomers. The observed negative crossing angle agrees with a right-handed GZ dimer, which has been proposed to correspond to the ligand-activated state. When the TYPE-C2 peptide was added to the system, Fscor ranges from 1.1 to 2.7. By contrast, for the N3-associated system the Fscor was low in all cases. These results suggest that the association of EphA2 TM dimers is reasonably strong when it is bound to TYPE7-C2, while the association is clearly much weaker when it interacts with TYPE-N3. A caveat of the interpretation of these Fscor parameters, is that they do not include the stabilizing interactions that the C2 peptide might contribute to stabilize the EphA2 dimer. Importantly, the crossing angle of the EphA2 TMD is similar in the presence and absence of TYPE-C2 and the distance between the two EphA2 TM peptides is only slightly increased, suggesting that TYPE-C2 binding does not cause a large conformational change.

We performed next simulations for a 2:2 stoichiometry, by adding a second copy of the TYPEs to the simulation. We observed similar results to those with the 2:1 stoichiometry (**Fig. S9**). In the EphA2:C2 systems, also performed in quadruplicate, the C2 peptides form homodimers in most of the simulations (1^st^ to 3^rd^). The C2 peptides associate with the EphA2 dimer to form a stable tetramer complex (**Fig. S9 A-D**). On the other hand, in the EphA2:N3 systems, the N3 peptides form a stable homodimer with EphA2 TM added on the outside, which prevents the latter from associating with each other in most of the simulations (1^st^ to 3^rd^) (**Fig. S9 E-H**). These results suggest that the stoichiometry of the TYPE peptide is not a key factor to understand its interaction with EphA2. For simplicity, we continued analysis using the 2:1 systems.

### MD simulations identify that TYPE-C2 uses several helical interfaces to bind to EphA2

To resolve the atomistic details of the association of the TYPE peptides with the EphA2 TMs, the CG models of the central conformers of C2 and N3 (2:1) systems were converted to AT resolution, and subjected to further refinement over 50 nanoseconds. A total of 12 simulations were conducted (4 each for the C2- and N3-associated EphA2 dimers, and 4 for the control EphA2 model system), yielding similar results. Only one representative simulation for each is discussed further for clarity. Given the slow conformational transition in the bilayer, cluster analysis of both the C2 and N3 complexes provided several conformational states. With a backbone RMSD of 4.3 Å from the control EphA2 system (**Fig. S6A**), the EphA2 dimer from the C2 system shows a helix crossing angle of -53° (**Fig. S6B**) and Fscor of 1.02. The EphA2 dimer from the N3 system shows a backbone RMSD of 6.3 Å from the control EphA2 system and an Fscor of 0, and the EphA2 helices remain separated by the peptide (**Fig. 6C**).

**Figure 6.**
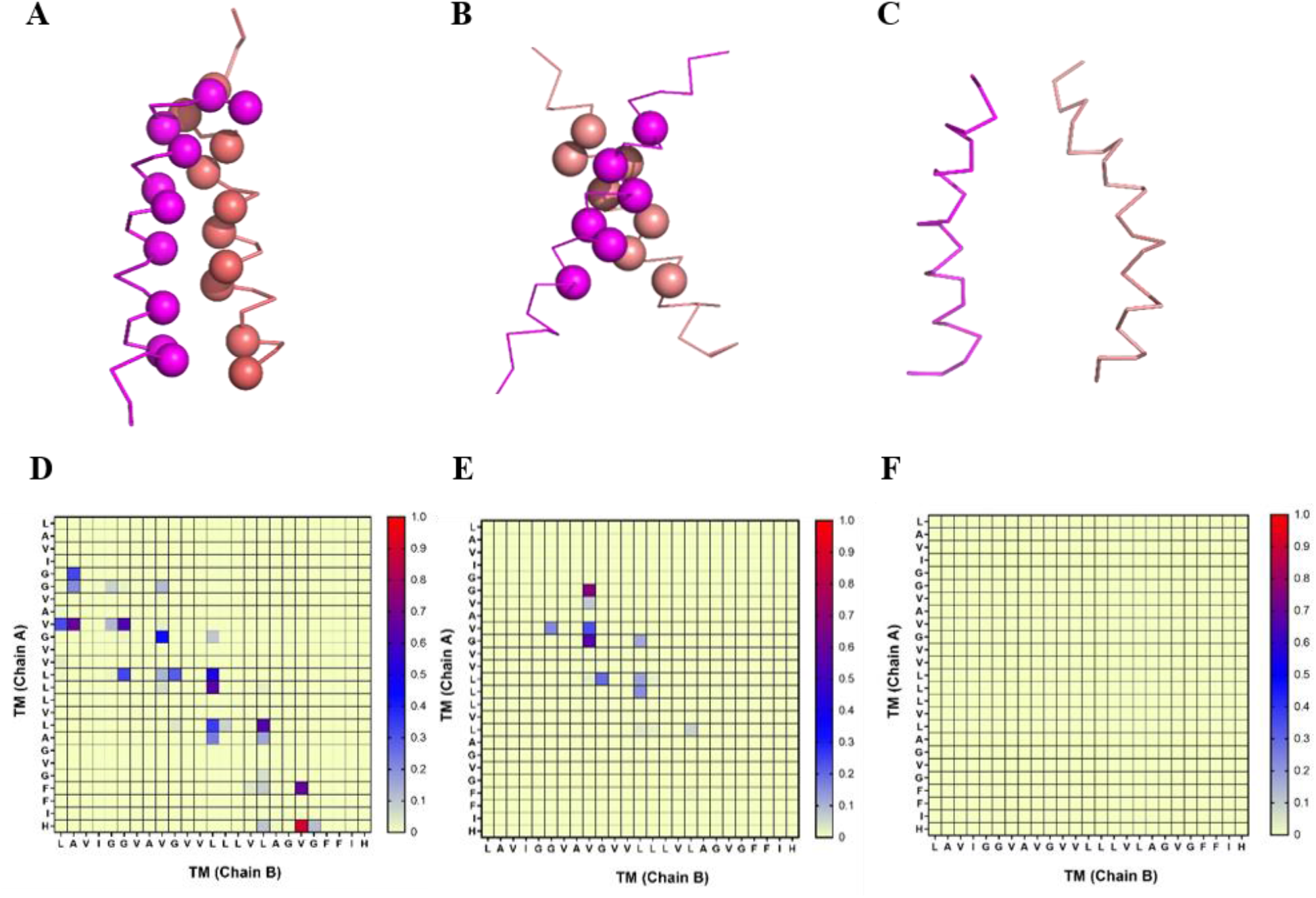
Comparison of EphA2 TM dimer Interfaces. ***Top* (A)** EphA2 TM dimer from the most populated conformer (helices shown in salmon for chain A and magenta for chain B). The Cα of residues found at the interaction interface are represented as spheres. **(B)** EphA2 TM dimer extracted from the most populated conformer of the EphA2:TYPE-C2 complex. **(C)** EphA2 TM dimer extracted from the most populated conformer of the EphA2:TYPE-N3 complex showing no EphA2 association. The TYPE7 and N3 peptides are not shown for clarity. ***Bottom***. Contact maps for the EphA2 TM dimer interface calculated with a cut off 4Å. The color scale (yellow to blue to red) indicates the fractional occupation of EphA2 contacts (0 to 1). **(D)** Contact map for the EphA2 TM dimer interface from the EphA2 model showing interaction throughout the TM interface. **(E)** The GZ motif is mainly involved in the interface extracted from the C2 system. **(F)** No contact interface is observed between the EphA2 TM dimers extracted from the EphA2:N3 system.

**Fig. 6** shows the contact map for the EphA2 TM interface in the presence of the C2 and N3 peptides compared to the isolated EphA2 TM dimer. The interaction interface of the EphA2 TMs shows the contact through the extended glycine zipper motif (A^536^, G^540^ and G^544^), in good agreement with the BiFC data (Fig. 4). In the case of the C2-associated system, the EphA2 TMs also associate *via* GZ residues (G^540^ and G^544^), but without involving A^536^, probably due to the slightly larger crossing angle. In contrast, there was no association between the EphA2 TMs in the case of the N3 complex.

Analysis of the interaction between TYPE-C2 and the two EphA2 helices allows generating a map with the residues that mediate the hetero-interaction (**Fig. 7**). We observed that TYPE-C2 binds largely to the center and the C_t_ end of helix A of EphA2, while binding to helix B is restricted to the N_t_ (**Fig. 7 A-C**). This asymmetric binding mode implies that TYPE-C2 uses different regions to interact with the two EphA2 helices. Additionally, it also indicates that the there are two helical interfaces, not one, that stabilize formation of the trimeric complex. Specifically, TYPE-C2 binds to helix B using the intact GZ interface of TYPE-C2. Helix B engages with the peptide using both GZ and interface 3 residues. The binding mode of TYPE-C2 to helix A of EphA2 was surprising, as it barely uses the predicted interfaces. Instead, the strongest interactions were observed between two C_t_ residues of TYPE-C2, as F^556^ promiscuously interacted with several C_t_ residues of EphA2, and F^557^ interacted with the last hydrophobic residue. Additionally, there was an electrostatic interaction at the C_t_ between the last E residue in TYPE-C2 and H^559^ of EphA2. Another important interaction was observed between one of the protonated Glu residues in C2 and the EphA2 TM residues. As shown in **Fig. S10A**, E^548^ of the C2 peptide initially (10 ns) interacts with the backbone CO of the G^540^ residue of chain A of EphA2, to then switch and interact with the backbone NH of the G^544^ of the chain B of EphA2 from 10 to 35 ns (**Fig. S10, B-C**). The N3 peptide (chain C) makes a parallel association with the separated chains A & B, and, again, uses different interfaces for both receptor helices. Additionally, the protonated E^545^ of the N3 peptide interacts with the backbone CO of V^537^ of the chain B of EphA2 throughout the AT simulation (**Fig. S10, D-F**) and it transiently interacts with the backbone NH of the V^541^ of chain B (**Fig. S10, E**). The TYPE-N3 peptide is noticeably tilted to the membrane normal, allowing such interactions.

**Figure 7.**
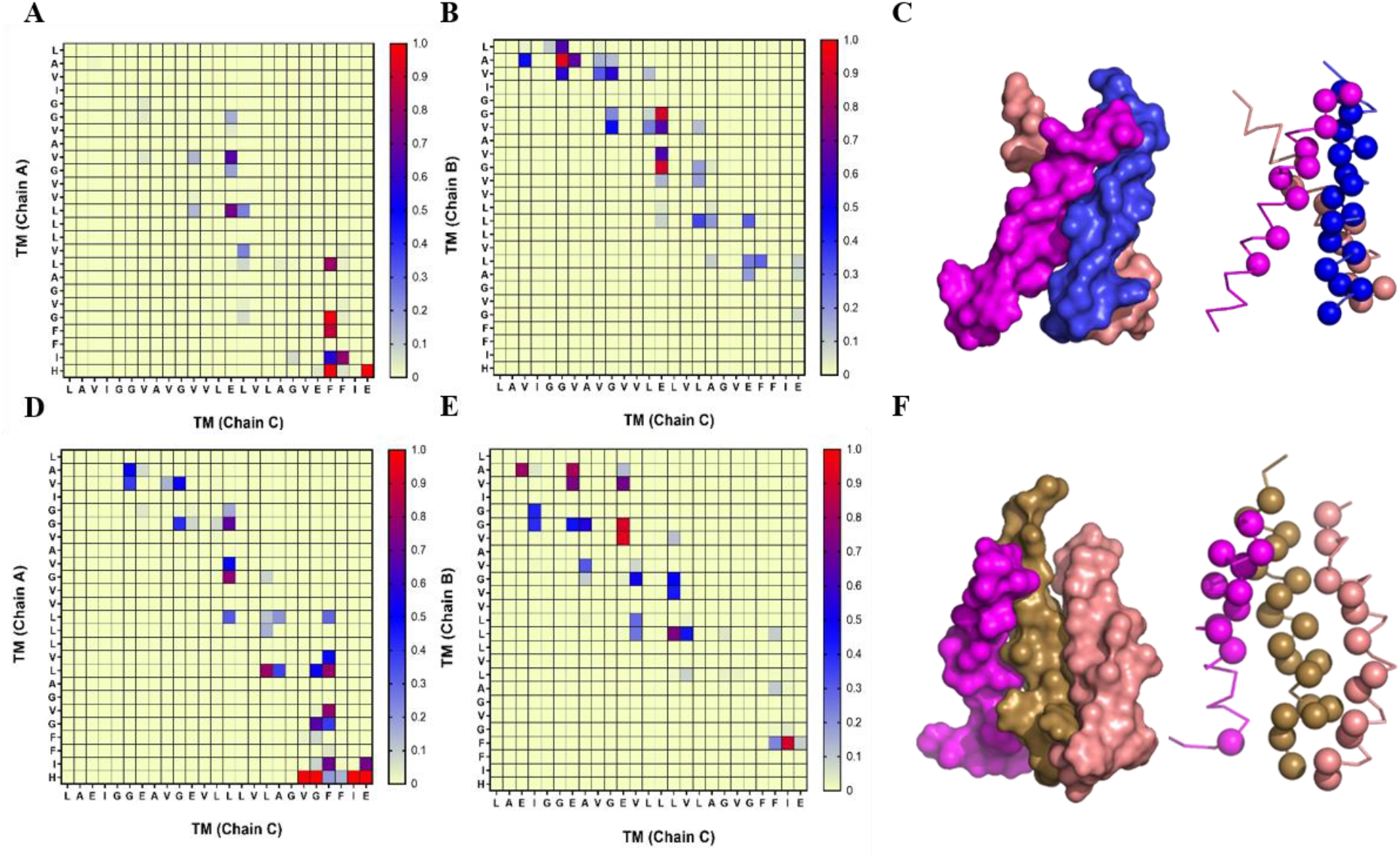
Contact map for the interface between the EphA2 TMs and the C2 or N3 peptides. **(A-B)** Contact interface between TYPE-C2 (chain C) and the EphA2 TMs (chains A and B). Same color scheme as Fig. 6 was used. **(C)** Surface (left) and sphere (right) representations of binding of TYPE-C2 peptide (blue) with the EphA2 TMs (magenta and salmon). Spheres mark the residues found at an interaction surface. **(D-E)** Contact interface of the N3 peptide (chain C) with EphA2 TMs (chains A and B). **(F)** Surface and sphere representations of N3 (shown in brown) binding to the EphA2 TMs. The Cα of residues found at the interaction interface are represented as spheres.

## Discussion

Membrane receptors operate as sensors that inform the cell of changes in their environment. Receptors are generally activated by binding of a ligand molecule present at the extracellular side of the membrane. Ligand binding triggers a conformational change in the receptor molecule, often involving dimerization [23], which activates the receptor. However, the structural and dynamic changes resulting from the activation event are poorly understood in most cases. In the receptor tyrosine kinase family, ligand binding induces a change in the ECR. This extracellular rearrangement is “ felt” by the TM, which is the domain tasked with transducing the signal into the cytoplasm. TM changes cause activation of the kinase domain at the ICR, which carries out its catalytic activity by phosphorylating tyrosine residues placed at different positions of the ICR. These P-Tyr residues trigger cytoplasmic signal transduction events. However, we are only beginning to understand how all these molecular events are coordinated, to create the adequate output that allows the cell to respond to the extracellular signals. An area particularly obscure is understanding the conformational changes that involve the TM segment.

We have previously described that the TYPE-C2 (TYPE7) peptide binds to the membrane region of EphA2 to activate this receptor [8]. TYPE7 is a conditional TM ligand that was created by strategically introducing E residues into the isolated TM&JM sequence of EphA2. The E residues were placed so they fall in adjacent helical turns. This staggered location is expected to prevent helix formation only at neutral pH, due to electrostatic repulsion between the aligned negatively charged side chains. A drop in pH will protonate E into the non-charged state. As a result, the overall peptide hydrophobicity increases to promote membrane insertion. This novel peptide design strategy builds on the rational design of the ATRAM peptide [24, 25], which also inserts into membranes upon a pH decrease due to the protonation of the E side chains. We previously showed that the TYPE-C2 peptide was able to bind to the membrane region of EphA2. However, the mechanism that TYPE-C2 uses to bind to and activate EphA2 is unknown.

In this work we tested the interaction with EphA2 of six TYPE peptides. The variants differ in that the E residues were placed at different positions along the TM helix. We hypothesized that these residues would act as a steric “ shield” and block *native* interactions at the helical surface where they align, while leaving the rest of the helix amenable to engage with the TM of EphA2. When we investigated the interaction of the TYPE peptides with the isolated TM domain of EphA2 using a reconstituted *in vitro* system, we observed that the C1 and C2 peptides caused large pH_50_ changes, suggesting that only these interactions are robust. We next studied the effect of the peptides on full-length EphA2 and assessed the functional relevance of the binding *via* a cellular motility assay, to test the ability of the peptides to bind to EphA2 and activate the receptor. The C1 and C2 TYPE peptides caused a strong reduction in cell migration, in agreement with the liposome results. Taken together, our data indicate that these two peptides contain intact the TM interfaces that allow for binding and receptor activation, even in the absence of ephrin ligand. These results additionally indicate that the interactions between the tail of E residues present at the C_t_ of all peptides (**Fig. 1A**) [8] are not enough to activate EphA2.

Binding of TYPE-C2 to EphA2 blocks the signaling cues that EphA2 provides to direct cells to migrate (**Fig. 3**). The TYPE-C2 peptide largely recapitulates the effect caused by binding of the ligand ephrinA1 to EphA2. When the ephrinA1 present at the plasma membrane of opposing cells engages with the ECR of EphA2, it causes the receptor to dimerize and undergo a conformational change that leads to activation of the intracellular kinase domain. In energetic terms, this involves that the presence of ephrinA1 bound to the LBD overcomes the free energy threshold needed to induce the dimerization and conformational change that activates the receptor. The monomer-dimer equilibrium free energy of EphA2 has been estimated to be ∼5 kcal/mole [5]. The free energy provided by binding of the two activating TYPE peptides with EphA2, are therefore expected to be larger than the stability difference between the inactive and active conformations of the receptor. A portion of this energetic contribution might be provided by the hydrogen bond that the residue E548 stablishes with the EphA2 peptide backbone **(Figs. 7 and S10)**. However, the stability provided by hydrogen bonds from polar residues in TM domains is typically small (0.5-1.0 kcal/mole in most cases) [26]. As a result, the main energetic contribution to TYPE7 binding are expected to be van der Waals interactions between hydrophobic residues at the TM helices. However, it has been previously observed that the presence of polar residues can promote the formation of hetero-trimers (see [27] and references therein). Additionally, acidic residues such as E can provide specificity to TM interactions [28-30], so the importance of polar interactions for the formation of the active complex should not be discounted.

To understand how the peptides activate EphA2, first we need to consider the conformation that the TMD of the receptor adopts in the active state, and in particular the interface that mediates TM dimerization. Sharonov *et al*. performed mutational studies of the HR and GZ interfaces of EphA2 and measured the activation of the receptor in cells [4]. Their data suggest that mutations at the GZ interface reduced EphA2 activity. However, these studies did not directly measure dimerization. In fact, studies of the effect of TM mutations on dimerization have not been performed in cells, to the best of our knowledge. Furthermore, to understand the effect of the TYPE peptides in our reconstituted system one must also know if the lack of the ICR and ECR interactions can cause a conformational change in the TMD dimer. This effect would be expected if, for example, the conformation of the receptor dimer was determined by interactions at the soluble domains, not the TMD. To evaluate this possibility, we measured the dimerization in cellular membranes of the isolated TM sequence of EphA2 (**Fig. 4**). The BiFC data clearly showed that the GZ interface mediates dimerization of the isolated TMJM_563_-EphA2 sequence, as it has been proposed for the full-length receptor. These data suggest an important piece of information to understand EphA2 dimerization: interactions at the ECR’s Cys-rich domain and LBD and the ICR, while able to modulate the strength of the dimerization [31, 32], do not qualitatively influence the conformation of the receptor. Taken together, these observations suggest that the TMD is a key domain defining the active conformation of EphA2. The good correspondence of the MD results with the effects caused by the TYPE-C2 peptide indicates that activation of EphA2 requires multiple inter-chain interactions that occur simultaneously in the membrane. The C1 and C2 peptides contain intact the GZ interface used for EphA2 to dimerize. As a result, it could theoretically compete for dimerization with an EphA2 monomer. This effect would reduce the levels of EphA2 dimer and as a result inhibit the receptor.

However, the cell migration data indicate that TYPE-C2 instead promotes dimerization. We propose this occurs because the EphA2 TM contains several dimerization interfaces. This allows TYPE-C2 to initially bind using another available interface, like residues in Interface 3. Once this interaction has been established, TYPE7 might engage laterally with the GZ dimer interface of EphA2 without breaking it. We propose that the formation of other interactions, including hydrogen bond formation at E548, occur at a later point to accommodate the hydrogen bond potential of the E side chain.

The conformation that TYPE7 adopts in the hetero-complex draws parallels to how a biological ligand activates another RTK. Specifically, the platelet-derived growth factor β receptor (PDGFβR) can be activated by E5, a TM protein of only 44 residues from papillomaviruses [33]. E5 binds to the TM of PDGFβR when it is in the dimer state [34]. Two copies of the dimeric E5 bind to PDGFβR, and clamp the active conformation of the receptor, driving signaling by the receptor [35]. The similarities with our binding model are striking, as one of the E5 helices interacts primarily with the N_t_ of one PDGFβR helix, and with the C_t_ of the other. Polar residues form hydrogen bonds with the TM of the receptor as well.

TYPE7 activates EphA2 with high specificity [8]. Other specific TM ligands included the traptamers developed by the DiMaio group, like LIL, which also promotes PDGFβR oligomerization in the active conformation [36]. The formation of supramolecular TM complexes where a peptide binds to multiple surfaces of the EphA2 TMD offers practical advantages as a therapeutic approach. An interaction with a single TMD would offer a small binding surface, where a limited number of stabilizing contacts could be stablished. On the other hand, a TMD dimer offers a significantly larger binding area, and importantly the crevice between the helices constitutes a long binding pocket where more stable contacts could be formed. This allows the C1 and C2 peptides to form a molecular clamp that stabilizes the active conformation of EphA2. Our data suggest that molecular clamping by membrane ligands can serve as a general mechanism to stabilize RTK into the active conformation, as a means to trigger receptor signaling.

## Methods

### Fluorescence labeling of peptides

Peptides were synthesized using F-moc chemistry, and purified to >95% purity, as assessed by HPLC and MALDI-TOF. For fluorescence experiments, the peptides were labeled at their N-terminus with an amine-reactive environmentally sensitive fluorescent reporter, Succinimidyl 6-(N-(7-nitrobenz-2-oxa-1,3-diazol-4-yl)amino)hexanoate (NBD-X, SE) (AnaSpec, Fremont, CA) [37]. For the reaction, peptides were weighed and dissolved in 0.1 M sodium phosphate buffer at pH ∼8, NBD-X, SE was weighed and dissolved in dimethylformamide (DMF) (Fisher Chemical, Thermo Fisher Scientific). Volumes were adjusted such that the reaction concentrations of peptide and dye were 1.6 x 10^−4^ M and 1.7 x 10^−3^ M, respectively, and the mole ratio was 11:1 dye:peptide. The reaction occurred for 2 hours gently shaking at room temperature in the dark. Excess dye was removed with a PD-10 desalting column from General Electric (Boston, MA). Unlabeled peptide was removed with HPLC on an Agilent (Santa Clara, CA) 1100 series HPLC with an Agilent Zorbax 300SB-C18 semi-preparative column. Peptides were separated by a linear gradient from 5–100% methanol in water. Both solvents contained 0.05% trifluoroacetic acid (TFA) (Sigma-Aldrich, St. Louis, MO). Fractions containing labeled peptide were concentrated by a rotary evaporator (Büchi Rotovapor-R, Flawil, Switzerland) to ∼10% of the original volume. Samples were then diluted to ≥ 90% water, aliquoted, frozen, and lyophilized (Labconco FreeZone 6, Fort Scott, KS) overnight. Labeling was confirmed by MALDI-TOF. Briefly, peptides were added to a saturated solution of α-cyano-4-hydroxy-cinnamic acid (TCI America, Portland, OR) in 70% acetonitrile (Fisher Chemical, Thermo Fisher) + 0.1 % TFA. The resulting solution was spotted onto a target plate and measured in a Bruker (Billerica, MA) MicroFlex mass spectrometer in positive mode.

### Fluorescence pH_50_ assay

Lyophilized, labeled peptides were resuspended in 10 mM sodium phosphate buffer and the concentration was determined by absorbance on an Agilent (Santa Clara, CA) Cary 100 UV-Vis spectrophotometer at 487 nm with a molar extinction coefficient of 18,482 M^-1^ cm^-1^. POPC was dried, resuspended, and extruded. For TM-EphA2 proteo-liposomes preparation, POPC and TM-EphA2 were dried together at a molar ratio of 500:1 of POPC:TMJM-EphA2_563_. Resuspension and extrusion were unchanged. Labeled peptides were incubated with liposomes in a “ master mix” for ≥ 1 hour. Samples were then added to a black 96-well plate and measured for fluorescence in a Biotek Cytation V plate reader. For this experiment, we found that a four-fold excess of TM-EphA2 over TYPE is important to observing interactions. The POPC:TYPE ratio is thus 2000:1. The high relative lipid concentration imparts noise in the fluorescence spectra. We found that the fluorescence center of mass is least sensitive to this noise, as compared to the spectral maximum or fluorescence intensity. For this reason, we collected spectra instead of endpoint fluorescence reads. Samples were excited at 470 nm and read from 500 to 600 nm with 1 nm step sizes a bandwidth of 9 nm. The gain was scaled to the brightest sample, and 50 measurements were collected per data point. The center of mass of the resulting fluorescence spectrum (520-600 nm) was calculated using equation 1,

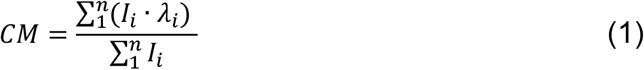

where I_i_ is the fluorescence intensity at wavelength λ_i_. The resulting centers of mass were plotted against the pH of the sample and fitted to equation 2 to determine the pH_50_ if two criteria were met: 1) the standard error of the pH_50_ value was above 0.5 pH units, and 2) the slope of the transition was less than 10.

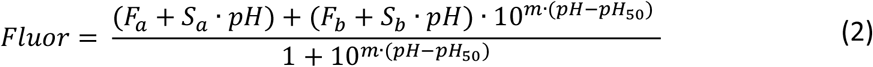

where F_a_ and F_b_ are the acidic and basic baselines, S_a_ and S_b_ are the slopes of the acidic and basic baselines, and m and pH_50_ are the slope and midpoint of the sigmoid, respectively. Circular dichroism pH_50_ values were obtained using the same equation to measure changes in mean residue ellipticity.

### Boyden chamber cell migration assay

1.1 × 10^6^ A375 cells were plated on a 10 cm plate 48 hours before plating chambers, and the plate was starved of serum 24 hours before plating. At zero hours, 600 μL of 10% FBS in DMEM containing treatment (0.5 μg/mL EphrinA1-Fc or 2 μM peptide) was placed below a 6.5 mm transwell polycarbonate membrane insert with an 8 μm pore size (Corning 3422). The 10 cm plate of serum starved cells at about 60% confluency was briefly trypsinized and transferred into serum-free DMEM to a density of 2.4 × 10^5^ cells/mL. 100 μL of the cell mixture was plated on top of the transwell inserts and incubated at 37°C with 5% CO_2_ approximately 18 hours. Upon completion, the top of the membrane was gently scraped to remove cells, the membranes were washed with PBS^++^, and cells on the bottom of the membrane were fixed for 30 minutes with methanol. Fixed cells were then stained with hematoxylin and eosin, membranes were washed, and dried membranes were mounted to microscope slides. 4 to 8 images of each membrane were taken using a Gen5 plate reader with a 10X color brightfield objective. The number of cells in each image were counted using ImageJ, averaged, and normalized to the control condition.

### Bimolecular fluorescence complementation (BiFC)

The TM sequences of EphA2 WT and mutants were PCR amplified and sub-cloned into BIFC plasmids (pBIFC-VN155 [VN] and pBIFC-VC173 [VC] at the amino end of the Venus Fluorescent Protein by using *EcoRI* and *XhoI* restrictions sites. All plasmid sequences were obtained by DNA sequencing (Macrogen). HEK293T cells were used to express the plasmids. Cells were grown in Dulbecco’s modified Eagle medium (Gibco) supplemented with 10% fetal bovine serum in 24-well plates at 37°C, 5% CO_2_ containing 2×10^6^ cells/plate. After 24 h of growing, 250 ng of each plasmid encoding VN and VC were transfected using 4 μL of 1 μg/mL PEI (Merck) for each μg of DNA. Twenty-five ng of a plasmid encoding Renilla luciferase under CMV promoter (pRL-CMV, Promega) was co-transfected for normalization purposes. Renilla were measured using the Renilla Luciferase Fash Assay Kit (Thermo), according to manufacturer’s instructions. Measurements of luminescence and fluorescence were performed 24 h post-transfection using a Multimode Plate Reader Victor X3 (Perkin Elmer). Immuno-identification of the samples was done using α-c-Myc rabbit antibody and followed by a secondary HRP-conjugated α-rabbit antibody (Merck). α-Histone H3 rabbit antibody was used as housekeeping protein. Chemi-luminescence was visualized by an ImageQuant LAS 4000 (GE Healthcare).

#### Computational modeling of the peptides

The NMR structure of EphA2 TM dimer (PDB ID: 2K9Y) [9] was obtained from www.rcsb.org. The TM region and the N-terminal residues of EphA2 from G^531^-K^563^ was extracted from the NMR structure and the remaining modified C-terminal residues from C^564^WN^566^ were modeled as an extended conformation of amino acids (ϕ, Ψ= ±120°) in PyMOL (The PyMOL Molecular Graphics System, Version 2.4. Schrödinger, LLC). The monomer structures, including the helices for the TM region of the TYPE7/C2 peptide [E^523^FQTLSPEGSGNLAVIGGVAVGVVL**E**LVLAGV**E**FFI**EEEEE**^563^] and the N3 peptide [E^523^FQTLSPEGSGNLA**E**IGG**E**AVG**E**VLLLVLAGVGFFI**EEEEE**^563^] were modeled as transmembrane helix from I^538^-I^558^ based on EphA2 TM NMR structure as the template using Modeller [38] and the remaining N-terminal residues (E^523^-V^537^ for C2 and E^523^-E^537^ for N3) and C-terminal residues (E^559^-E^563^ for both C2 and N3) were modeled as an extended conformation (ϕ, Ψ= ±120°) in PyMOL. The C2 peptide has 7 mutations and the N3 peptide has 8 mutations as shown in **Fig. S1**.

### Coarse-grain (CG) molecular dynamics simulation

To check the influence of the C2/N3 peptides on the dimerization of EphA2 TMs, we built an initial all-atom model system configuration that contains two EphA2 TM monomers and the C2 or the N3 peptide, with all three peptide units placed perpendicular to the membrane and 50 Å apart from each other. As a reference for the above system, we also ran simulations with the same system set-up but omitting the C2/N3 peptide, i.e. the two EphA2 TM peptides by themselves as the control. To study if the C2/N3 peptides can form homo/heterodimers and how these peptides affect the dimerization of EphA2 TMs, we built another system with two EphA2 TM monomers and two C2- or N3 peptides again, with all four units placed perpendicular to the membrane and 50 Å apart from each other.

The atomistic (AT) modeled systems with EphA2 TMs: N3/C2 (2:1) and EphA2 TMs alone, each as placed as above, were converted to coarse-grained (CG) representation using the *martinize2*.*py* code (https://github.com/marrink-lab/vermouth-martinize) considering the secondary structure based on the DSSP assignment. The MARTINI 3 force field was released as a beta version in July of 2018, but has been subsequently modified [39]. CG simulations were performed using Gromacs version 2016.5.[40] The *insane*.*py* script [41] was used for setting up of the POPC bilayer (typically 678 lipid and 19,680 CG water molecules for 2:1 peptide systems; 306 lipids and 4,735 CG water molecules for control EphA2 TM dimer systems) around the peptides. The pH of the system was 4.5, setting all the Glu residues in the peptides and Histidine residues of EphA2 to be protonated. The systems were equilibrated for 500 ps. The electrostatic interactions were shifted to zero between 0 and 11 Å and the Lennard-Jones interactions where shifted to zero between 9 and 11 Å. The V-rescale thermostat was used with a reference temperature of 320 K in combination with a Berendsen barostat at 1 bar reference pressure, with a coupling constant of 1.0 ps, a compressibility of 3.0 × 10^−4^ bar ^-1^. The integration time step was 20 fs and all the simulations were run in quadruplicate for 4 μs.

### All-atom (AT) molecular dynamics simulation

Conformational clustering of all the coarse grain simulation trajectories was performed with a backbone RMSD cut-off 6 Å. Conformers, identified as the nearest to the center of the major cluster were selected for further analysis. These coarse grain conformations of the central structures were then converted to all atom resolution using the Backward tool of Martini [42]. All-atom conformations of the selected structures were further refined in a POPC bilayer for 50 ns each at 320 K by use of the gromos-96 43a1 force field in GROMACS 2016 [40]. The packing of lipid bilayers around the transmembrane regions was performed using the InflateGRO approach [43]. The final system was re-solvated with water molecules (SPC) and then neutralized by appropriate counter ions (Na^+^ and Cl^−^). All the systems were equilibrated twice, first for 100 ps under NVT followed by 1 ns under appropriate NPT conditions before the MD studies. Numerical integrations were performed with a step size of 2 fs and coordinates were updated every 0.2 ps. Bonds to hydrogens were constrained with LINCS with the order 4. The non-bonded pair list cut-off was 1.2 nm with a grid function. The particle mesh Ewald (PME) algorithm was implemented for long-range electrostatic calculations.

It should be noted that the results from such AT refinement need to be interpreted with caution, as Best and coworkers [44] have recently shown that the charmm36 potential function led to non-native energy barriers and also likely non-native intermediates on the associate pathway of glycophorin A, a model TM helix dimer peptide. However, here we used an older potential function gromos-96 43a1 and assume that the 50 ns simulation length is short enough to equilibrate the grown-in atoms upon CG to AT conversion but not to substantially alter the mode of helix-helix packing.

### Data Analysis

The PREDDIMER webserver was used for analyzing the TM dimers based on the Fscor and helix crossing angle [22]. Analysis of the trajectories was done by implementing the modules built into GROMACS. The contact maps for the TM regions between the helices were calculated with a cut off 4 Å for all the backbone and side-chain atoms. Sequence alignments of the EphA2 TM and the N3/C2 peptides were done using ClustalX [45]. Data were plotted in GraphPad Prism (version 6 for Windows, GraphPad Software, La Jolla California USA, www.graphpad.com). Inter-helical distances between the centers of mass of the TM regions were calculated and the clustering were performed using the module of the GROMACS.

## Acknowledgements

This work was supported by National Institutes of Health grants R01GM120642 (to F.N.B) and R01GM112491 (to M.B.), and PROMETEU2019/065 grant from Generalitat Valenciana and BFU2016-79487-P grant from the Spanish Ministry of Science and Universities (to I.M.). We thank Dr. Siewert Marrink (University of Groningen, Netherlands) in whose lab the MARTINI 3 force field was developed.

## References

[1] Barquilla A, Pasquale EB. Eph receptors and ephrins: therapeutic opportunities. Annu Rev Pharmacol Toxicol. 2015;55:465–87.

[2] Tandon M, Vemula SV, Mittal SK. Emerging strategies for EphA2 receptor targeting for cancer therapeutics. Expert Opin Ther Targets. 2011;15:31–51.

[3] Pasquale EB. Eph receptor signalling casts a wide net on cell behaviour. Nat Rev Mol Cell Biol. 2005;6:462–75.

[4] Sharonov GV, Bocharov EV, Kolosov PM, Astapova MV, Arseniev AS, Feofanov AV. Point mutations in dimerization motifs of the transmembrane domain stabilize active or inactive state of the EphA2 receptor tyrosine kinase. J Biol Chem. 2014;289:14955–64.

[5] Singh DR, Ahmed F, King C, Gupta N, Salotto M, Pasquale EB, et al. EphA2 Receptor Unliganded Dimers Suppress EphA2 Pro-tumorigenic Signaling. J Biol Chem. 2015;290:27271–9.

[6] Nikolov DB, Xu K, Himanen JP. Homotypic receptor-receptor interactions regulating Eph signaling. Cell Adh Migr. 2014;8:360–5.

[7] Himanen JP, Yermekbayeva L, Janes PW, Walker JR, Xu K, Atapattu L, et al. Architecture of Eph receptor clusters. Proc Natl Acad Sci U S A. 2010;107:10860–5.

[8] Alves DS, Westerfield JM, Shi X, Nguyen VP, Stefanski KM, Booth KR, et al. A novel pH-dependent membrane peptide that binds to EphA2 and inhibits cell migration. Elife. 2018;7.

[9] Bocharov EV, Mayzel ML, Volynsky PE, Mineev KS, Tkach EN, Ermolyuk YS, et al. Left-handed dimer of EphA2 transmembrane domain: Helix packing diversity among receptor tyrosine kinases. Biophys J. 2010;98:881–9.

[10] Stefanski KM, Russell CM, Westerfield JM, Lamichhane R, Barrera FN. PIP2 promotes conformation-specific dimerization of the EphA2 membrane region. J Biol Chem. 2020.

[11] Russ WP, Engelman DM. The GxxxG motif: a framework for transmembrane helix-helix association. JMolBiol. 2000;296:911–9.

[12] Anderson SM, Mueller BK, Lange EJ, Senes A. Combination of Calpha-H Hydrogen Bonds and van der Waals Packing Modulates the Stability of GxxxG-Mediated Dimers in Membranes. Journal of the American Chemical Society. 2017;139:15774–83.

[13] Lemmon MA, Flanagan JM, Hunt JF, Adair BD, Bormann BJ, Dempsey CE, et al. Glycophorin A dimerization is driven by specific interactions between transmembrane alpha-helices. J Biol Chem. 1992;267:7683–9.

[14] Orzaez M, Lukovic D, Abad C, Perez-Paya E, Mingarro I. Influence of hydrophobic matching on association of model transmembrane fragments containing a minimised glycophorin A dimerisation motif. FEBS Lett. 2005;579:1633–8.

[15] Scott HL, Westerfield JM, Barrera FN. Determination of the Membrane Translocation pK of the pH-Low Insertion Peptide. Biophys J. 2017;113:869–79.

[16] Miao H, Li D-Q, Mukherjee A, Guo H, Petty A, Cutter J, et al. EphA2 mediates ligand-dependent inhibition and ligand-independent promotion of cell migration and invasion via a reciprocal regulatory loop with Akt. Cancer Cell. 2009;16:9–20.

[17] Garcia-Murria MJ, Duart G, Grau B, Diaz-Beneitez E, Rodriguez D, Mingarro I, et al. Viral Bcl2s’ transmembrane domain interact with host Bcl2 proteins to control cellular apoptosis. Nat Commun. 2020;11:6056.

[18] Lucendo E, Sancho M, Lolicato F, Javanainen M, Kulig W, Leiva D, et al. Mcl-1 and Bok transmembrane domains: Unexpected players in the modulation of apoptosis. Proc Natl Acad Sci U S A. 2020;117:27980–8.

[19] Brayan Grau MJ, Maria Jesús García-Murria, Waldemar Kulig, Ilpo Vattulainen, Ismael Mingarro, Luis Martínez-Gil. The role of hydrophobic matching on transmembrane helix packing in cells. Cell Stress. 2017;1:90–106.

[20] Andreu-Fernandez V, Sancho M, Genoves A, Lucendo E, Todt F, Lauterwasser J, et al. Bax transmembrane domain interacts with prosurvival Bcl-2 proteins in biological membranes. Proc Natl Acad Sci U S A. 2017;114:310–5.

[21] Zhang L, Sodt AJ, Venable RM, Pastor RW, Buck M. Prediction, refinement, and persistency of transmembrane helix dimers in lipid bilayers using implicit and explicit solvent/lipid representations: microsecond molecular dynamics simulations of ErbB1/B2 and EphA1. Proteins. 2013;81:365–76.

[22] Polyansky AA, Chugunov AO, Volynsky PE, Krylov NA, Nolde DE, Efremov RG. PREDDIMER: a web server for prediction of transmembrane helical dimers. Bioinformatics. 2014;30:889–90.

[23] Westerfield JM, Barrera FN. Membrane receptor activation mechanisms and transmembrane peptide tools to elucidate them. J Biol Chem. 2020;295:1792–814.

[24] Nguyen VP, Palanikumar L, Kennel SJ, Alves DS, Ye Y, Wall JS, et al. Mechanistic insights into the pH-dependent membrane peptide ATRAM. J Control Release. 2019;298:142–53.

[25] Nguyen VP, Alves DS, Scott HL, Davis FL, Barrera FN. A Novel Soluble Peptide with pH-Responsive Membrane Insertion. Biochemistry. 2015;54:6567–75.

[26] Bowie JU. Membrane protein folding: how important are hydrogen bonds? CurrOpinStructBiol. 2011;21:42–9.

[27] Zhang L, Polyansky A, Buck M. Modeling transmembrane domain dimers/trimers of plexin receptors: implications for mechanisms of signal transmission across the membrane. PLoS One. 2015;10:e0121513.

[28] Gratkowski H, Lear JD, DeGrado WF. Polar side chains drive the association of model transmembrane peptides. ProcNatlAcadSciUSA. 2001;98:880–5.

[29] Zhou FX, Merianos HJ, Brunger AT, Engelman DM. Polar residues drive association of polyleucine transmembrane helices. Proc Natl Acad Sci U S A. 2001;98:2250–5.

[30] Bano-Polo M, Martinez-Gil L, Wallner B, Nieva JL, Elofsson A, Mingarro I. Charge pair interactions in transmembrane helices and turn propensity of the connecting sequence promote helical hairpin insertion. J Mol Biol. 2013;425:830–40.

[31] Shi X, Hapiak V, Zheng J, Muller-Greven J, Bowman D, Lingerak R, et al. A role of the SAM domain in EphA2 receptor activation. Sci Rep. 2017;7:45084.

[32] Singh DR, Ahmed F, Paul MD, Gedam M, Pasquale EB, Hristova K. The SAM domain inhibits EphA2 interactions in the plasma membrane. Biochim Biophys Acta. 2017;1864:31–8.

[33] Talbert-Slagle K, Marlatt S, Barrera FN, Khurana E, Oates J, Gerstein M, et al. Artificial transmembrane oncoproteins smaller than the bovine papillomavirus E5 protein redefine sequence requirements for activation of the platelet-derived growth factor beta receptor. J Virol. 2009;83:9773–85.

[34] Talbert-Slagle K, DiMaio D. The bovine papillomavirus E5 protein and the PDGF beta receptor: It takes two to tango. Virology. 2008.

[35] Karabadzhak AG, Petti LM, Barrera FN, Edwards APB, Moya-Rodriguez A, Polikanov YS, et al. Two transmembrane dimers of the bovine papillomavirus E5 oncoprotein clamp the PDGF beta receptor in an active dimeric conformation. Proc Natl Acad Sci U S A. 2017;114:E7262–E71.

[36] Heim EN, Marston JL, Federman RS, Edwards APB, Karabadzhak AG, Petti LM, et al. Biologically active LIL proteins built with minimal chemical diversity. Proc Natl Acad Sci U S A. 2015;112:E4717–25.

[37] Chattopadhyay A, London E. Spectroscopic and ionization properties of N-(7-nitrobenz-2-oxa-1,3-diazol-4-yl)-labeled lipids in model membranes. Biochim Biophys Acta. 1988;938:24–34.

[38] Sali A, Blundell TL. Comparative protein modelling by satisfaction of spatial restraints. J Mol Biol. 1993;234:779–815.

[39] Souza PCT AR, Barnoud J, Thallmair S, Faustino I, Grunewald F, Patmanidis I, Abdizadeh H, Bruiniks BMH, Wassenaar TA, Kroon PC, Melcr J, Nieto V, Corradi V, Khan HM, Domanski J, Javanainen M, Martinez-Seara H, Reuter N, Best RB, Vattulainen I, Monticelli L, Periole X, Tieleman P, de Vries AH, Marrink SJ. Martini 3: A general purpose force field for coarse-grain molecular dynamics Nature Methods. 2021;In Press.

[40] Abraham M, Murtola T, Schultz R, Szilard P, Smith JC, Hess B, et al. GROMACS: High performance molecular simulations through multi-level parallelism from laptops to supercomputers. Software X. 2015:19–25.

[41] Wassenaar TA, Ingolfsson HI, Bockmann RA, Tieleman DP, Marrink SJ. Computational Lipidomics with insane: A Versatile Tool for Generating Custom Membranes for Molecular Simulations. J Chem Theory Comput. 2015;11:2144–55.

[42] Wassenaar TA, Pluhackova K, Bockmann RA, Marrink SJ, Tieleman DP. Going Backward: A Flexible Geometric Approach to Reverse Transformation from Coarse Grained to Atomistic Models. J Chem Theory Comput. 2014;10:676–90.

[43] Kandt C, Ash WL, Tieleman DP. Setting up and running molecular dynamics simulations of membrane proteins. Methods. 2007;41:475–88.

[44] Domanski J, Sansom MSP, Stansfeld PJ, Best RB. Balancing Force Field Protein-Lipid Interactions To Capture Transmembrane Helix-Helix Association. J Chem Theory Comput. 2018;14:1706–15.

[45] Larkin MA, Blackshields G, Brown NP, Chenna R, McGettigan PA, McWilliam H, et al. Clustal W and Clustal X version 2.0. Bioinformatics. 2007;23:2947–8.

